# Coordination of rhythmic RNA synthesis and degradation orchestrates 24-hour and 12-hour RNA expression patterns in mouse fibroblasts

**DOI:** 10.1101/2023.07.26.550672

**Authors:** Benjamin A. Unruh, Douglas E. Weidemann, Shihoko Kojima

**Affiliations:** Department of Biological Sciences, Fralin Life Sciences Institute, Virginia Tech, Blacksburg, VA USA

**Keywords:** circadian rhythms, 4-thiouridine, rhythmic gene expression, RNA dynamics, metabolic labeling, mouse fibroblast

## Abstract

Circadian RNA expression is essential to ultimately regulate a plethora of downstream rhythmic biochemical, physiological, and behavioral processes. Both transcriptional and post transcriptional mechanisms are considered important to drive rhythmic RNA expression, however, the extent to which each regulatory process contributes to the rhythmic RNA expression remains controversial. To systematically address this, we monitored RNA dynamics using metabolic RNA labeling technology during a circadian cycle in mouse fibroblasts. We find that rhythmic RNA synthesis is the primary contributor of 24 hr RNA rhythms, while rhythmic degradation is more important for 12 hr RNA rhythms. These rhythms were predominantly regulated by *Bmal1* and/or the core clock mechanism, and interplay between rhythmic synthesis and degradation has a significant impact in shaping rhythmic RNA expression patterns. Interestingly, core clock RNAs are regulated by multiple rhythmic processes and have the highest amplitude of synthesis and degradation, presumably critical to sustain robust rhythmicity of cell-autonomous circadian rhythms. Our study yields invaluable insights into the temporal dynamics of both 24 hr and 12 hr RNA rhythms in mouse fibroblasts.

## Introduction

Circadian rhythms are the internal timing mechanisms that allow organisms to anticipate and respond to daily changes in the environment, and most organisms on Earth exhibit circadian rhythms in their behavior, physiology, and biochemical processes ^1, 2^. Identification of rhythmically expressed genes has been an active area of research in recent decades, as they are believed to ultimately drive the diverse range of rhythmic biological processes. Circadian transcriptome studies have demonstrated that rhythmic RNA expression is pervasive in various organisms and their rhythmic patterns must be tightly regulated to ensure correct period, phase, and amplitude (reviewed in ^3^).

For a molecule, such as RNA or protein, to exhibit rhythmic expression, its synthesis, degradation, or a combination of both must be rhythmic. In addition, the average half-life of the molecule must be short enough (< 10 hrs) ^4–6^. The mechanisms of driving rhythmic RNA synthesis (i.e., transcription) are well established in mice. The “core” mammalian clock mechanism, comprised of interlocking transcription-translation feedback loops, is considered the primary force of generating cell-autonomous circadian rhythms and rhythmic gene expression ^7^. Many of the ‘core’ clock genes, such as *Arntl* (or *Bmal1*), *Clock*, *Ror,* and *Nr1d* (or *Rev-Erb*), encode transcription factors and activate rhythmic RNA transcription by recognizing target DNA motifs in the promoter sequence of the target genes^7, 8^. These studies uncovered that approximately 5% to 40% of transcriptomes are rhythmic in mouse depending on the tissue and more than 50% of transcripts are rhythmic in at least one tissue^9, 10, 11, 12^. Previous studies estimated the extent of rhythmic RNA synthesis in driving rhythmic RNA expression, however, there has been significant variation between studies, ranging from 20 to 85 % in mouse liver^5, 13–15^.

Compared to RNA synthesis, however, much less is known about how rhythmic RNA degradation is regulated and how much rhythmic RNA degradation contributes to driving and sustaining rhythmic RNA expression. Several ‘post-transcriptional’ mechanisms (e.g., miRNA, poly(A) tail length, RNA methylation, and RNA-binding protein, etc) regulate rhythmic RNA expression and many of them directly influence mRNA stability^16–28^. This suggests that rhythmic RNA degradation is an important post-transcriptional mechanism driving rhythmic RNA expression. In support of this, RNA degradation of core clock genes, such as *Period2* (*Per2*) and *Cryptochrome1* (*Cry1*), is faster when their level is declining compared to when it is accumulating^21, 22^. A recent study combining transcriptomic data with mathematical estimation also predicted that at least 35% of rhythmically expressed mRNA are rhythmically degraded^5^.

Given that rhythmic degradation is energetically cost-effective to drive RNA rhythms^5^ and the regulatory mechanism for RNA degradation is more diverse^29^, we hypothesized that rhythmic RNA degradation is one of the major post-transcriptional mechanisms regulating rhythmic RNA expression. To test this, we analyzed RNA dynamics using metabolic labeling approach throughout the circadian cycle on a genome-wide scale in NIH3T3 cells. Deciphering the dynamics of rhythmic RNA expression deepens our understandings of how the core circadian clock machinery orchestrates RNA rhythms and ultimately regulates rhythmic downstream processes.

## Results

### Monitoring rhythmic RNA dynamics in mouse fibroblasts using metabolic labeling

To monitor RNA dynamics over a circadian cycle, we used metabolic labeling approach, as it allows us to simultaneously measure the synthesis and degradation kinetics from a single sample, and minimally interferes with cell metabolism and physiology ^29, 30^. We used *Bmal1-luc* cell line, in which a luciferase reporter gene driven by *Bmal1* promoter is stably expressed in NIH3T3 cells ^31^, allowing us to monitor the rhythmicity of the cells in real-time. We also knocked-down *Bmal1,* one of the core clock genes critical for cell-autonomous rhythmicity ^32^, to evaluate whether the rhythmicity is driven by *Bmal1* or the core clock machinery (Fig. 1A). We synchronized cells by adding 50% horse serum to the culture media ^33^ and labeled newly synthesized RNAs with 4-thiouridine (4sU) every two hours for two hours starting at T22 (i.e., 22hrs after the end of serum shock) (Fig. 1A). We opted to use ‘pulse-in’ procedure, as this caused little or no disruption to cellular rhythmicity (Fig. S1A). We extracted total RNA at 12 time points from T24 to T46, corresponding to one circadian cycle (Fig. 1A). The 4sU-labeled RNAs were subsequently biotinylated and pulled-down with a pull-down efficiency of 50-75 % in all samples (Fig. S1B), comparable to other reports ^34, 35^. We used higher time resolution (2 hr interval) with less biological replicas (n=1) given the cost, potential technical variability between batches, and most importantly the community guidelines for genome-scale analysis of biological rhythms ^36^. We then subjected both total and newly synthesized RNA fractions to transcriptomic analyses (RNA-seq and 4sU-seq, respectively) (Fig. 1A). As expected, reads from RNA-seq were mapped predominantly to exons, while those from 4sU-seq were mapped more to introns and intergenic regions both in control and *Bmal1* KD cells (Fig. 1B). We used INSPEcT ^37^, an R package, to derive the rates of RNA synthesis, processing, and degradation, and to quantify the levels of pre- and mature RNAs (Fig. 1C) at each time point, as it was built for a dynamic system and hence can handle time-series data, and can computationally normalize the pull down efficiency between each time point ^37^.

**Figure 1.**
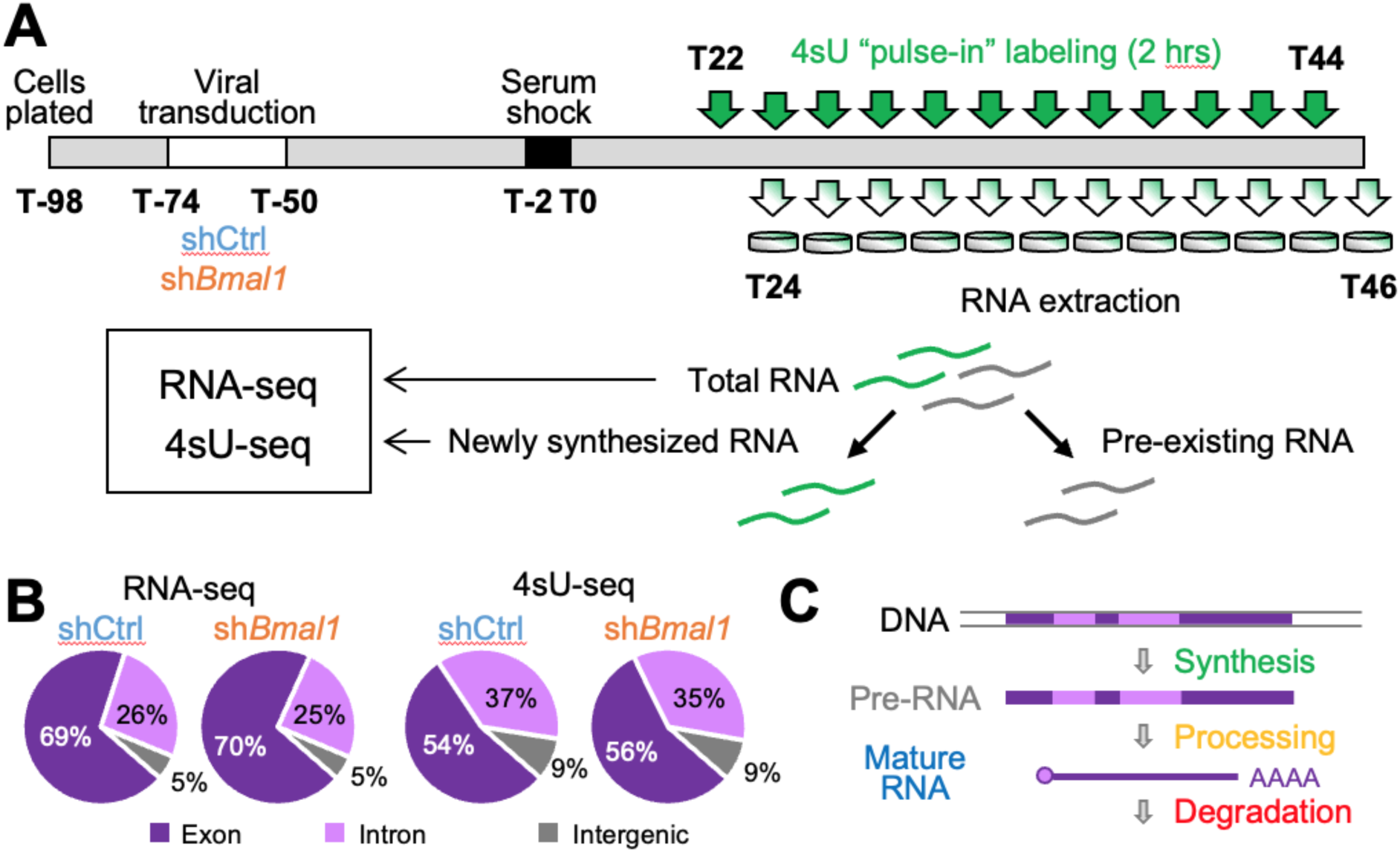
Rhythmic 4sU-seq and RNA-seq analysis in NIH3T3 cells. (A) Experimental design. Cells were plated 24 hrs before lentiviral transduction (72 hrs) containing scrambled shRNA (shCtrl) or shRNA targeting *Bmal1* (sh*Bmal1*). Cells were then incubated with 50% horse serum for 2 hrs to synchronize cellular circadian rhythms. 22 hours after the serum shock, 400mM 4sU were added to the culture media for 2 hrs to label newly synthesized RNAs of the first sample. At the end of the 2hr incubation, RNAs were harvested. We repeated this procedure every 2 hrs starting from T22 until T44. Newly synthesized RNAs were then isolated from total RNAs, and both total and newly synthesized RNAs were subjected to RNA-seq analysis. (B) Percentage of read counts mapped to exon (dark purple), intron (light purple), or intergenic region (grey). The average from all time points in each dataset was used. (C) Schematic representation of the lifecycle of an RNA.

We first performed extensive quality checks of our dataset. First, successful knock-down (KD) of *Bmal1* was verified both by *Bmal1-luc* bioluminescence output and the levels of *Bmal1* RNA in both RNA-seq and 4sU-seq (Fig. S1C-E). Second, we verified that the RNA expression level was very strongly correlated between RNA-seq and 4sU-seq data at each time point (Pearson correlation coefficient > 0.95) in all samples (Fig. S2). Third, the level of mature RNA correlated with the RNA synthesis and degradation rates, either positively or negatively, as expected (Fig. S3). Fourth, we compared our dataset to the previous 4sU-seq study in NIH3T3 cells ^38^ and found a strong correlation for mature RNA level (Pearson r = 0.672) and synthesis rate (Pearson r = 0.578) between the two datasets (Fig. S4A-B). The correlation for RNA half-lives was still strong (Pearson r = 0.611), although the median half-life of our dataset was shorter than that of the previous report (this study: 2.27 hr, Schwanhäusser: 10.04 hr) (Fig. S3C-D). This could be due to: 1) the difference between static (Schwanhäusser) vs dynamic systems (this study), 2) the effect of serum shock in our dataset, or 3) the effect of viral transduction in our dataset, although absolute RNA half-lives are often inconsistent between datasets even with the same method and the same cells^29^. Nevertheless, the distribution of RNA half-lives in our dataset was similar to what was estimated from mouse liver over a circadian cycle and free of serum shock or viral transduction ^39^. Overall, these data demonstrate that our datasets are of high quality.

### Bmal1 or the core clock machinery drives rhythmic RNA synthesis but not degradation

sAfter the analysis with INSPEcT, we obtained a total of 11313 RNAs that met our analytical criteria. We then used MetaCycle ^40^ (20 < τ < 28) to statistically determine rhythmicity of the rates of RNA synthesis, processing, and degradation, and the levels of pre- and mature RNAs. We did not use a stringent statistical threshold to determine the rhythmicity, because circadian transcriptome output in cell cultures is not as robust as those from tissues ^41^. We identified 653, 272, 336, 685, and 6 RNAs whose synthesis rate, pre-RNA level, processing rate, mature RNA level, and degradation rate, respectively, was rhythmic in control cells (meta2d B.H.q < 0.25) (Fig. 2A, Datafile 1). The number of rhythmic mature RNAs was comparable with a previous study using NIH3T3 cells with a similar statistical threshold and experimental design (Fig. S1F). Rhythmic mature RNAs also included most of the core clock genes, such as *Arntl* (or *Bmal1*), *Period* (*Per*)*1-3*, *Nr1d1-2* (or *Rev-erbα/²*), and *Dbp* (Datafile 1). Globally, the rhythmic mature RNA expression had a bimodal phase distribution and was highly enriched around T26 and T38 (Fig. 2B-C). A similar phase distribution pattern was observed for pre-RNAs (Fig. 2B-C). Both RNA synthesis and processing had a single peak at around T38 (Fig. 2B-C). Nevertheless, the biological significance of rhythmic RNA processing is unclear, as processing includes multiple steps, such as (potentially co-transcriptional) splicing, capping, polyadenylation, and nuclear export, and the processing time is much shorter than a circadian cycle (median: 12.8 min in control cells) (Fig. 2H: blue). Rhythmic degradation of RNAs was not as robust in *Bmal1-luc* NIH3T3 cells (Fig. 2A-C), compared to previous theoretical studies^5, 6^.

**Figure 2.**
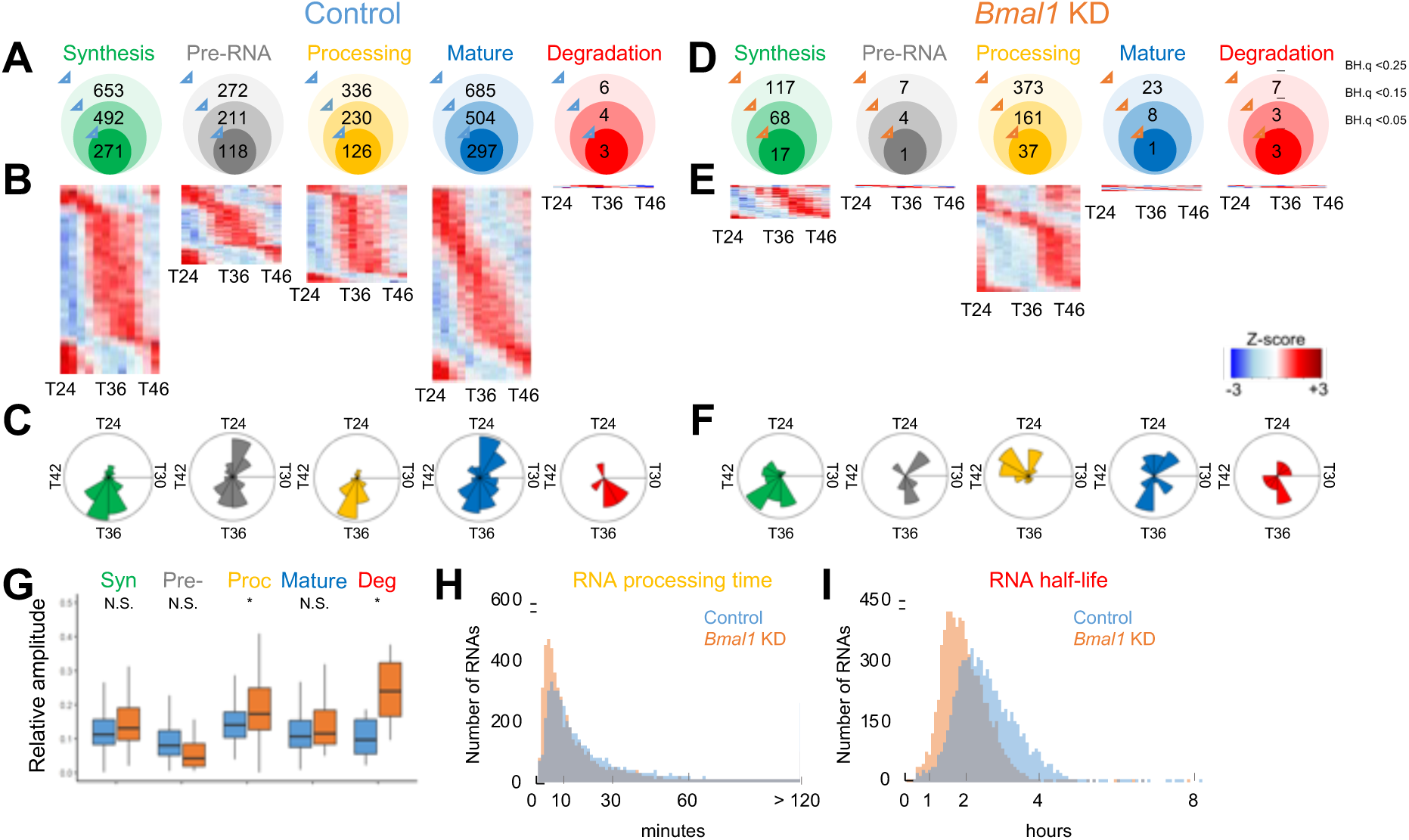
Rhythmicity of RNA dynamics in NIH3T3 cells. A total of 11313 genes were analyzed in all datasets. (A, D) Number of RNAs whose RNA synthesis (green), processing (yellow), or degradation as well as unprocessed (grey) or mature RNA (blue) levels was rhythmic in control (A) or *Bmal1* KD (D) cells with various statistical thresholds (meta2d_BH.q < 0.25, 0.15, or 0.05). (B, E) Phase-sorted heatmap of RNAs with rhythmic RNA synthesis (green), unprocessed RNA levels (grey), processing (yellow), mature RNA levels (blue), and degradation (red) in control (B) or *Bmal1* KD (E) cells with the rhythmicity cutoff as metaCycle_BH.q < 0.25. Color represents Z-scores, in which red is high while blue is low. Each line represents one RNA. (C, F) Circular histograms representing the peak phase distribution of rhythmic RNA synthesis (green), processing (yellow), or degradation as well as pre (grey) or mature RNA (blue) levels. Each bin represents 2 hrs. Radius line represents number of genes at each tick mark, and the most outer line is 170 (synthesis), 60 (pre-RNA), 120 (processing), 120 (mature RNA), and 3 (degradation) RNAs in control and 30 (synthesis), 3 (pre-RNA), 100 (processing), 6 (mature RNA), and 3 (degradation) RNAs in *Bmal1* KD cells. (G) Distribution of relative amplitude (meta2d_rAMP) for rhythmic RNAs in control (blue) and *Bmal1* KD (orange) cells. Boxplots represent two quartiles ± 1.5 interquartile range from median (midline). *; p < 0.05, N.S.; p > 0.05 (two-tailed Student’s t-test). (H, I) Distribution of RNA processing time (H) and half-life (I) in control (blue) and *Bmal1* KD (orange) cells. The average of all time points was used.

We also analyzed the effect of *Bmal1* and found that the number of rhythmic RNAs and processes were significantly lower in *Bmal1* KD cells. We identified 117, 7, 373, 23, and 7 RNAs whose synthesis rate, pre-RNA level, processing rate, mature RNA level, and degradation rate, respectively, was rhythmic in *Bmal1* KD cells (meta2d B.H.q < 0.25) (Fig. 2D, Datafile 2). The phase distribution patterns were similar between control and *Bmal1* KD cells, except for processing (Fig. 2C, F). Nevertheless, there was little or no overlap between control and *Bmal1* KD cells in any of the processes or RNA levels, and some RNAs even gained rhythmicity in *Bmal1* KD cells (Datafile 1, 2). In fact, relative amplitudes (rAMPs) of RNA synthesis, pre-RNA, and mature RNA levels were comparable between control and *Bmal1* KD cells, while that of processing and degradation were higher in *Bmal1* KD cells (Fig. 2G). Unexpectedly, both processing time (median WT: 12.8 min, KD: 8.7 min, p = 2.2*10^-16^ two-sided Mann-Whitney U test) and RNA half-life (median WT: 1.99 hr, KD: 1.44 hr, p = 2.2*10^-16^ two-sided Mann-Whitney U test) were shorter in *Bmal1* KD cells (Fig. 2H-I), indicating that *Bmal1* and/or the core clock machinery regulates RNA processing and degradation either directly or indirectly. Overall, these data indicate that *Bmal1* or the core clock machinery drives rhythmic RNA synthesis but not degradation.

### Rhythmic RNA expression is primarily regulated by rhythmic RNA synthesis in NIH3T3 cells

We next focused on the 685 rhythmic mature RNAs (Fig. 2A-C) and analyzed which of the three processes contribute most to the rhythmicity of their mature RNA levels. Because the venn diagram method underestimates the number of rhythmic transcripts overlapping in different datasets ^42^, and all the current statistical algorithms to compare rhythmicity are applicable only to transcriptomic data and not between RNA levels and process rates (DODR ^39^, LimoRhyde 1/2 ^43, 44^, and CompareRhythms ^42^), we instead calculated adjusted B.H.q values for rhythmicity of process rates using only the set of 685 rhythmic mature RNAs^43, 44^ (Datafile 3).

By using the adjusted B.H.q < 0.25 as a new statistical threshold, we found 402 RNAs had at least one rhythmic process (Fig. 3A). The most significant contribution was rhythmic RNA synthesis, as 389 (57%) mature RNAs had rhythmic RNA synthesis (Fig. 3A-B). Both processing and degradation only had a modest contribution (Fig. 3A-B). The rhythmicity of both mature RNA levels and RNA synthesis were significantly dampened in *Bmal1* KD cells, although not completely abolished (Fig. 3B-D, Datafile 3), supporting the role of *Bmal1* as a transcriptional activator. A principal component analysis demonstrated that consecutive samples for RNA synthesis, but not processing nor degradation, were placed in a spiral configuration in control, but not in *Bmal1* KD cells (Fig. 3C). These data indicate that RNA synthesis is the primary contributor of RNA rhythms and *Bmal1* or the core clock machinery plays a major role in driving rhythmic RNA synthesis.

**Figure 3.**
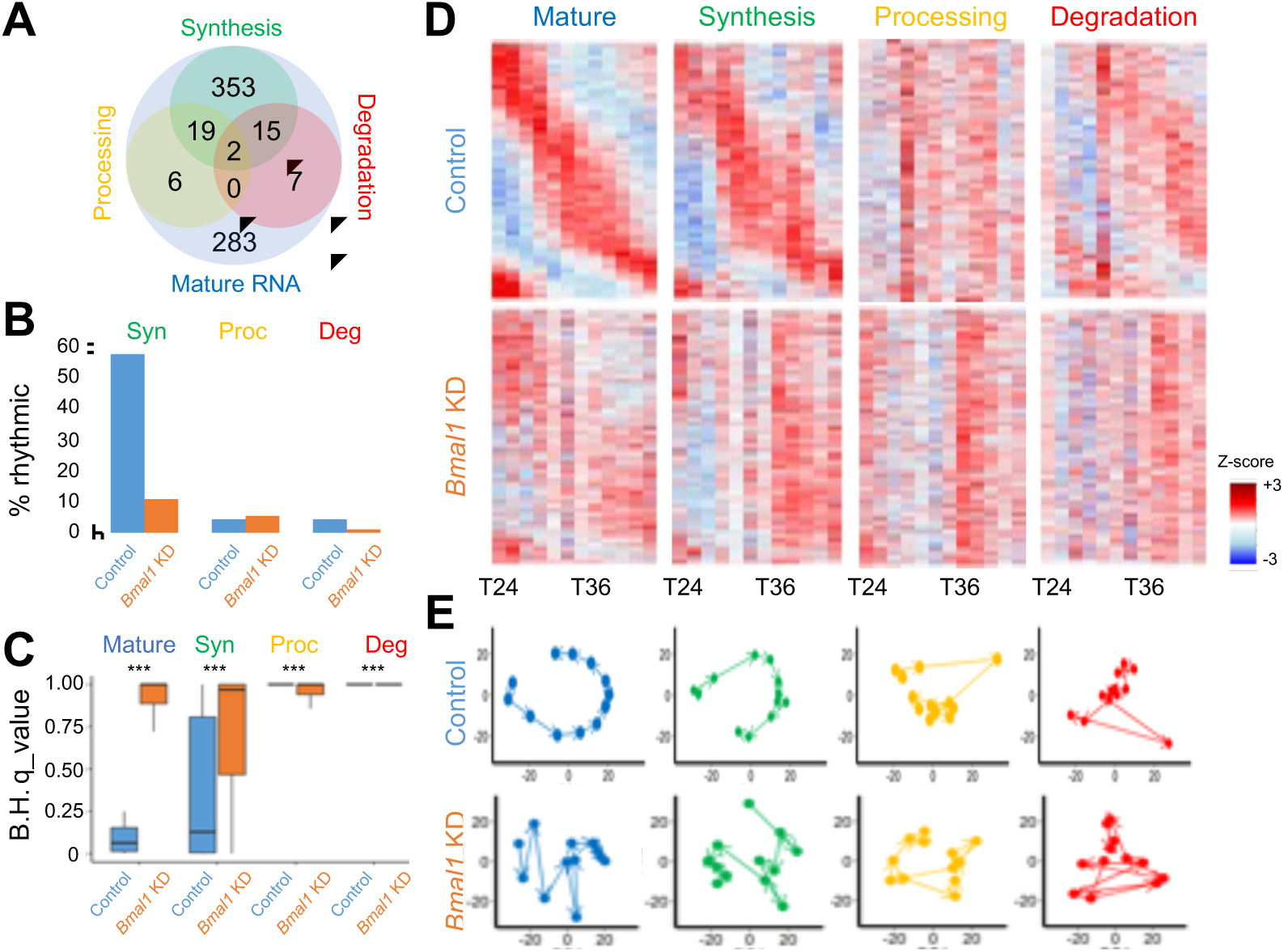
Rhythmic RNA synthesis is the major driver for rhythmic RNA expression in NIH3T3 cells. 685 rhythmic mature RNAs in control cells were analyzed in all datasets. (A) Venn diagram showing the number of RNAs whose RNA synthesis (green), processing (yellow), and degradation (red) was rhythmic (metaCycle_BH.q < 0.25). (B) The percentage of RNAs with statistically significant synthesis, processing, or degradation rhythms (MetaCycle BH.q < 0.25) among the 685 rhythmic mature RNAs in control cells. (C) Distribution of adjusted B.H.q values for the 685 rhythmic RNAs in each dataset. Box plots represent two quartiles ± 1.5 interquartile range from median (midline). ***; p < 1.0 x 10^13^ (two - tailed Mann-Whiney U test). (D) Phase sorted heatmap of 685 rhythmic mature RNAs (blue) for their RNA synthesis (green), processing (yellow), and degradation (red) in control (top) and *Bmal1* KD (bottom) cells. Color represents Z-scores, in which red is high while blue is low. Each line represents one RNA. (E) Principal component analyses of the 685 rhythmic RNAs in each dataset in control (top) or *Bmal1* KD (bottom) cells. The first two components in each dataset are plotted against each other. Arrows represent the progression of time from T24 to T46. Variation of each principal components in control cells; Mature RNA (PC1 – 49%, PC2 – 29%), Synthesis (PC1 – 40%, PC2 – 22%), Processing (PC1 – 28%, PC2 – 16%), and Degradation (PC1 – 23%, PC2 – 18%) or in *Bmal1* KD cells; Mature RNA (PC1 – 39%, PC2 – 24%), Synthesis (PC1 – 38%, PC2 – 26%), Processing (PC1 – 33%, PC2 – 16%), and Degradation (PC1 – 33%, PC2 – 24%).

### The effect of interplay between rhythmic synthesis and degradation on RNA rhythms

Theoretical studies predicted how RNA dynamics can provide flexibility on both the phase and amplitude of rhythmic RNA expression ^5, 6, 45^. Since our dataset provides additional information for RNA dynamics that cannot be retrieved from RNA-seq or 4sU-seq data alone, they present an ideal opportunity to test these predictions. To this end, we again focused on the 685 rhythmic mature RNAs (Fig. 3) and used meta2d_phase and meta2d_rAMP values, even though some were not statistically rhythmic (meta2d B.H.q > 0.25).

The peak phase of rhythmic mature RNA expression was predicted to be anywhere throughout the 24 hr cycle, when the amplitude of RNA degradation rhythms is greater than that of RNA synthesis rhythms ^6^. This is largely true, as while 71% of rhythmic RNAs peaked within -3/+3 hours of the peak RNA synthesis when the amplitude of synthesis rate is higher than that of degradation rate, the RNA peak phase was more widely distributed when the amplitude of degradation rate was higher than that of synthesis rate (p = 5.33 x 10^-8^, Watson-Wheeler test) (Fig. 4A). The peak phases of mature RNA were also predicted to be within 6 hrs of that of RNA synthesis if the RNA synthesis is rhythmic but degradation is constant ^6, 45^. We only found one RNA that matches this prediction, in which both RNA synthesis and degradation were rhythmic and had more than 6 hrs of phase difference between RNA synthesis and mature RNA level (*Abcd4*: ϕλPhase [mat-syn] = -10.08 hr). When RNA degradation is rhythmic (Fig. 4B: red), the phase difference between mature RNA levels and RNA synthesis were clustered into two time zones: close to 0 hr (cluster 1) or between -6 to -9 hrs (cluster 2), whereas 65% of rhythmic RNAs had their phase peak within -1 and + 6 hrs from that of RNA synthesis when RNA synthesis was rhythmic (Fig. 4B: green). Interestingly, all but one in cluster 1 had their RNA synthesis rhythmic (Datafile 3), while none in cluster 2 had rhythmic RNA synthesis. We also found that the effect of RNA half-lives on the phase difference between RNA synthesis and mature RNA levels was minimal (Fig. 4C), presumably because half-lives of all RNAs were less than 10 hrs in our dataset (Fig. 2I).

**Figure 4.**
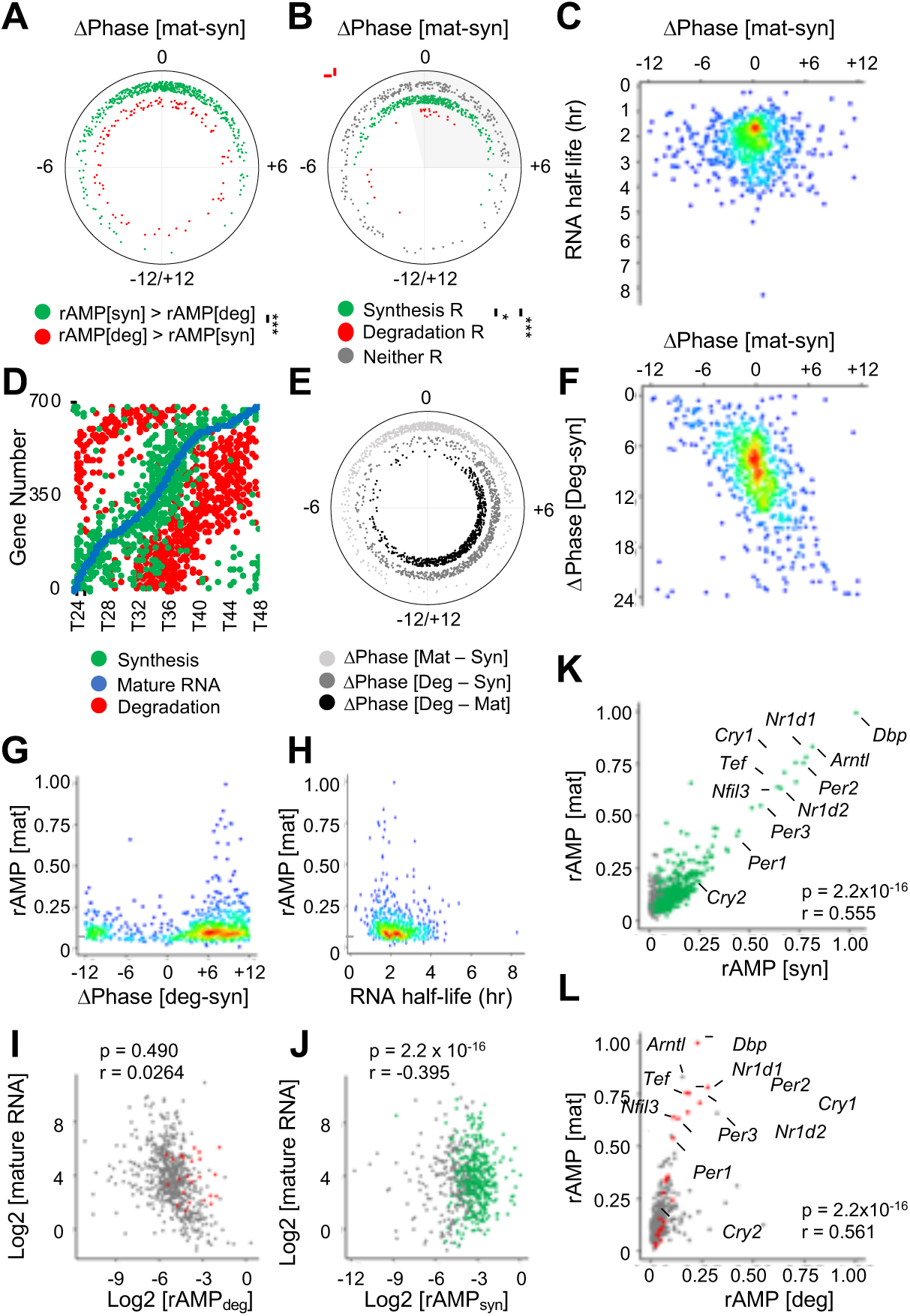
Effects of rhythmic RNA synthesis and degradation on the phase and amplitude of RNA rhythms. 685 rhythmic mature RNAs in control cells were analyzed in all datasets. (A, B) Phase difference between mature RNA expression and RNA synthesis (DPhase [mat-syn] in hr) when the relative amplitude of RNA synthesis (rAMP [syn]) is higher than that of RNA degradation (rAMP[deg]) (green) or the relative amplitude of RNA degradation (rAMP [deg]) is higher than that of RNA synthesis (rAMP[syn]) (red) (A) or when RNA synthesis is rhythmic (green), degradation is rhythmic (red), and neither is rhythmic (grey) (meta2d B.H. q-value < 0.25) (B). ***; p < 0.001, *; p < 0.05 (Watson-Wheeler test). (C) Relationship between RNA half-lives (average of all time points) and phase difference between mature RNA expression and RNA synthesis (DPhase [mat-syn]). (D) Peak phases of RNA synthesis (green), mature RNA (blue), and degradation (red). (E) Phase difference (hr) between mature RNA levels and RNA synthesis (light grey), mature RNA and RNA degradation (dark grey), and RNA synthesis and degradation (black). (F) Relationship between the phase difference between mature RNA levels and RNA synthesis (DPhase [mat-syn]) and the phase difference between RNA synthesis and degradation (DPhase [deg-syn]). (G, H) Relationship between the relative amplitude of mature RNA expression (rAMP [mat]) and the phase difference between RNA synthesis and degradation (DPhase [deg-syn]) (G) or RNA half-lives (H). (I, J) Correlation between the level of mature RNA and relative amplitude of RNA degradation rate (rAMP [deg]) (I) or synthesis rate (rAMP [syn]) (J). All values are log2 transformed. (K, L) Correlation between the relative amplitude of mature RNA (rAMP [mat]) and that of RNA degradation rate (K) or synthesis rate (L). (I – L) r: Pearson correlation coefficient. Colored dots represent those that are statistically rhythmic (meta2d_B.H. q-value <0.25) for synthesis (J, K) or degradation (I, L) whereas grey dots are statistically not rhythmic.

We also noticed from the heatmap (Fig. 3D) that RNA degradation appeared to have a rhythmic pattern in control cells, even though the amplitude is low and most RNA does not reach statistical threshold for their rhythmicity (Fig. 3B-D). Interestingly, the peak phase of RNA degradation was approximately 6.02 hrs (median) delayed from that of RNA synthesis while the phase difference between RNA synthesis and mature RNA level was only 0.10 hr (median) (Fig. 4D-F). The anti-phasic relationship between synthesis and degradation was predicted to lead to the highest amplitude^6^; however, in our study, the highest amplitude of RNA rhythm was achieved when the phase difference between RNA synthesis and degradation was 8.52 hrs (*Dbp*: meta2d_phase_[syn]_ = 22.56 and meta2d_phase_[deg]_ = 7.08) and a majority of rhythmic RNAs had their relative amplitude around the median (= 0.11) (Fig. 4G: grey dotted line). Higher relative amplitude of mature RNA levels was also observed when RNA half-lives were shorter in general as predicted ^6^, although the RNA half-life of the highest amplitude RNA (*Dbp*: 2.17 hrs) was not the shortest among all (Fig. 4H). Because the RNA half-life is significantly shorter and its range is narrower than previously reported^38, 46–49^, the effect of half-lives on the phase or amplitude of rhythmic RNA expression is minimal in our dataset.

Contrary to the theoretical prediction, an increased amplitude of RNA degradation did not lead to higher mean RNA levels, even though RNA half-lives are less than ten hours ^6^. Instead, we found the amplitude of RNA degradation negatively correlated with the mean RNA levels (Fig. 4I). The amplitude of RNA synthesis did not have any impact on the level of mean RNA levels, either (Fig. 4J). Regardless, the amplitude of RNA rhythm is higher when that of synthesis or degradation is also higher (Fig. 4K-L), as predicted ^6^. Most remarkably, most of the core clock genes with rhythmic RNA expression had high amplitude in RNA synthesis and/or degradation (Fig. 4K-L), presumably critical for sustaining robust rhythmic RNA expression. Overall, our data provided experimental support for some of the theoretical predictions, but not all, suggesting that some parameters are biologically constrained.

### Most core clock genes are regulated by multiple rhythmic processes

Some RNAs were regulated by more than one rhythmic process, and 15, 19, and 2 RNAs were regulated by a combination of rhythmic synthesis and degradation, synthesis and processing, or all three processes, respectively (Fig. 3A). Interestingly, most core-clock genes were regulated by more than one rhythmic process (Fig. 5, Datafile 3), suggesting that multiple layers of regulation are necessary to sustain robust rhythmic RNA expression of core clock RNAs.

**Figure 5.**
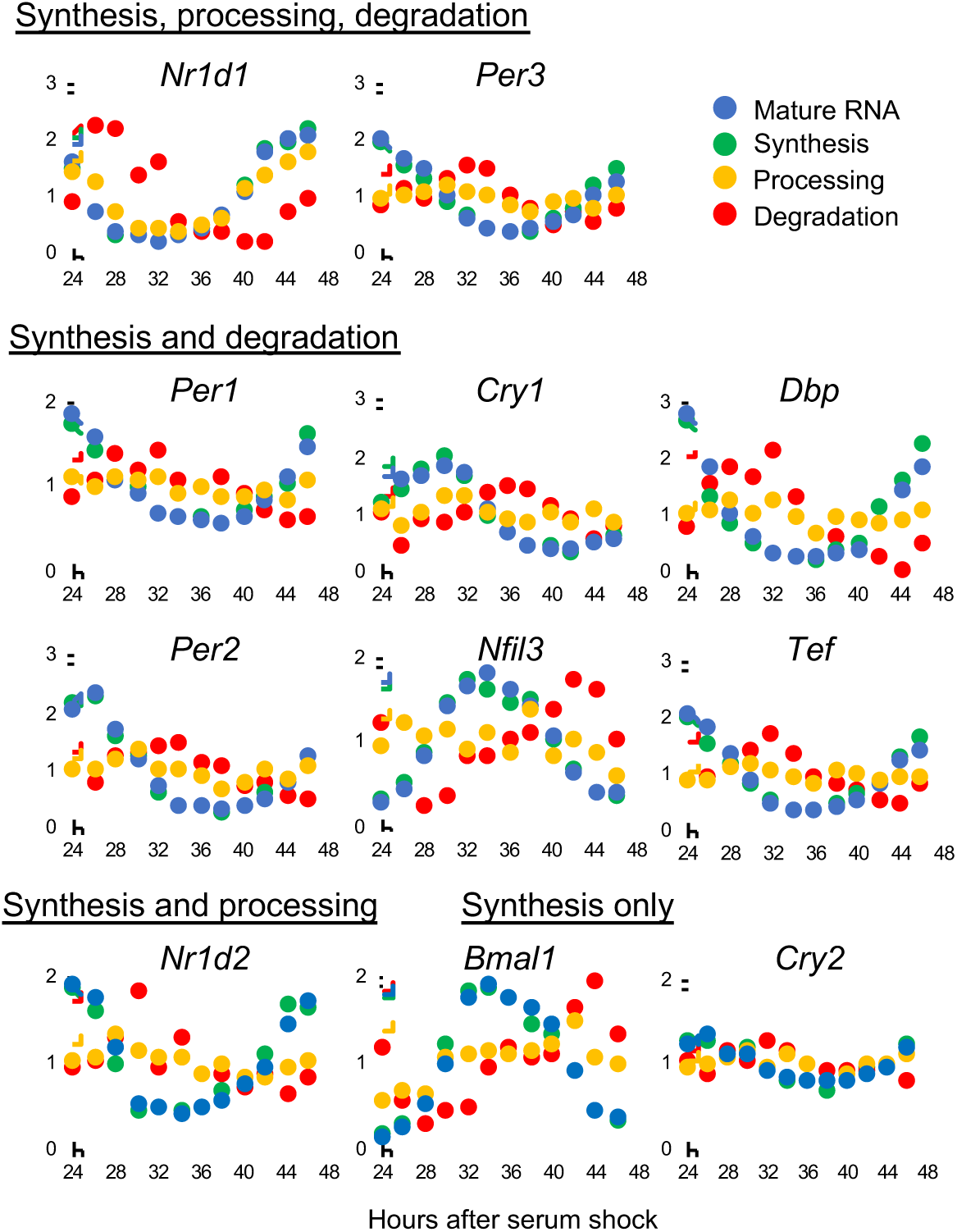
Multiple rhythmic processes regulate most core clock gene expression. Time series data of mature RNA levels (blue), RNA synthesis rate (green), processing rate (yellow), and degradation rate (red) in control cells. All values were normalized by dividing values at each time point by the average across all time points for each gene. Note that *Clock*, *Npas2*, and *Rora* were expressed but not rhythmically. *Rorc* was not detected.

Puzzlingly, some RNAs did not have any rhythmic processes that could account for the rhythmicity of the mature RNA levels (Fig. 3A). These RNAs are generally less robust and had low amplitude (Fig. S5A-B). Since the threshold to define rhythmicity of synthesis and degradation was arbitrary (meta2d B.H.q <0.25), we tested whether all the RNA rhythms could be explained by at least one rhythmic process by setting a different statistical threshold for rhythmicity. However, some rhythmic RNAs have adjusted B.H. q-value = 1 for all three processes (Datafile 3). Conversely, several RNAs had rhythmic processes, but not rhythmic mature RNAs (Datafile 1) and in fact, 35 RNAs had two processes being rhythmic (synthesis and processing) and 345 and 280 RNAs had one process being rhythmic (synthesis and processing, respectively) without RNA rhythms (Datafiles 1). Their arrhythmic RNA expression cannot be explained solely by their RNA half-lives, because the median RNA half-lives of non rhythmic RNAs are shorter than that of rhythmic RNAs and they are also comparable regardless of whether one or more processes are rhythmic or not (Fig. S5C). Given that our statistical threshold for rhythmicity is not too stringent (meta2d B.H.q <0.25), some of the rhythms (or lack thereof) may be due to statistical noise or false positives. In addition, the rhythmicity mismatch between RNA levels and kinetic processes may derive from a limitation in statistically comparing the rhythmicity between different types of datasets.

### 12hr RNA rhythms are primarily regulated by rhythmic RNA degradation in NIH3T3 cells

In addition to ∼ 24 hr RNA rhythms, ultradian RNA rhythms with the period of approximately 12 hrs or 8 hrs have been observed both in mouse tissues and cultured cells^41, 50–52^ and are considered important for metabolism and stress response to dawn/dusk transition^41, 50–52^. The regulatory mechanisms driving ultradian gene expression, however, have remained largely elusive. Using our transcriptomic dataset with high temporal resolution, we analyzed RNA dynamics of ultradian RNA expression. By pre-assigning the period range in MetaCycle, we identified 161 and 293 mature RNAs with the period of approximately 12 (10 < τ < 14) or 8 (6.7 < τ < 9.3) hrs, respectively (meta2d B.H.q < 0.25) (Datafiles 4-5). This number is most likely underestimated, as our method precludes RNA rhythms superimposed by those with different periods including 24 hrs^51, 52^. These 8 hr- or 12 hr-RNA rhythms were not as robust as 24 hr RNA rhythms (Fig. 6A-B), and the median RNA half-lives of 8 hr- and 12 hr-rhythmic RNAs were slightly shorter than that of 24 hr rhythmic RNAs (Fig. 6C) although their distribution was not that much different from each other (Fig. 6D).

**Figure 6.**
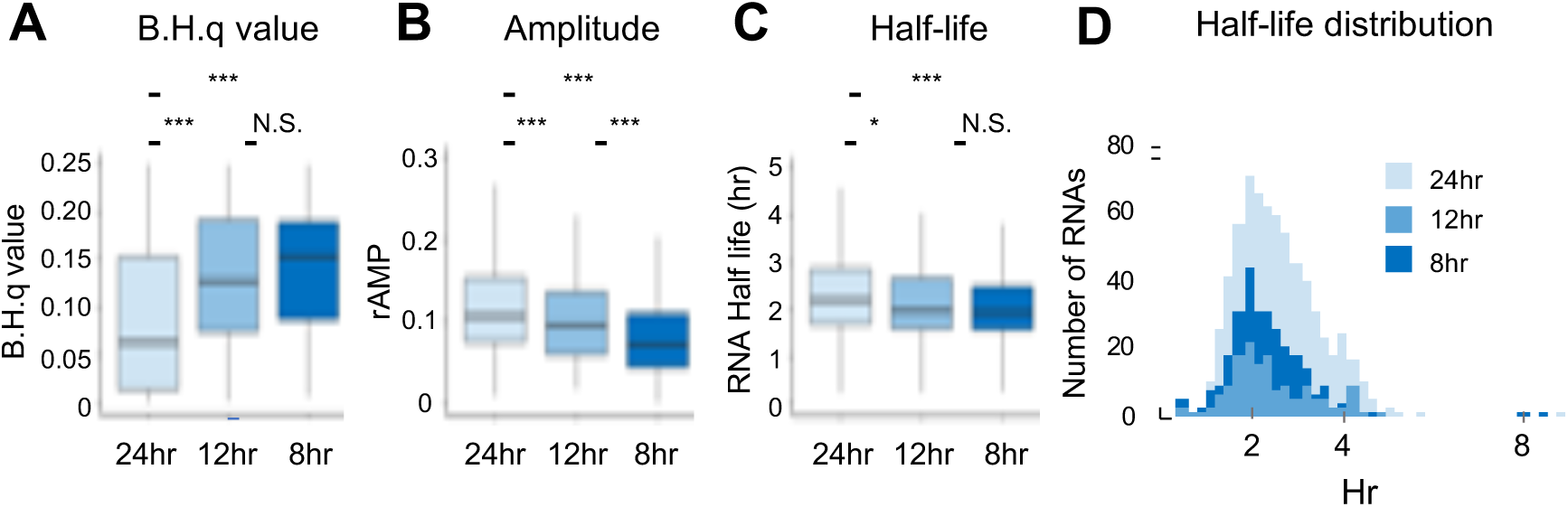
Comparison between 8hr, 12hr and 24 hr RNA rhythms. (A) Distribution of B.H.q values, (B) relative amplitude (meta2d_rAMP), and (C) RNA half-lives (hr) of RNAs with 24 hr, 12 hr, and 8 hr rhythms (meta2d B.H.q < 0.25). Box plots represents two quartiles ± 1.5 interquartile range from median (midline). ***; p < 0.001, *; p < 0.05 (A: two-tailed Mann Whitney U Test. B and C: two-tailed Students’ t-test). N.S.; p > 0.05. (D) Distribution of RNA half-life for rhythmic RNAs with 24 hr (median: 2.20 hr), 12 hr (median: 1.97 hr), and 8 hr (median: 1.94 hr) periods.

We also detected hundreds of RNAs undergoing rhythmic synthesis, processing, and degradation with a period of 12 hrs in control cells (Fig. S6A). Interestingly, 12 hr rhythms of RNA synthesis and mature RNA levels were dampened in *Bmal1* KD cells (Fig. S6B, Datafiles 4, 6). This is a stark contrast to mouse liver and mouse embryonic fibroblasts (MEFs), in which *Bmal1* or the core circadian clock mechanisms have little or no effect on the 12 hr RNA rhythms^52^. *Xbp1s*, a basic region leucine zipper transcription factor, is expressed with the ∼12 hr rhythmicity and predominantly regulates 12 hr gene expression in mouse liver or MEFs ^52^. Curiously, however, the expression of *Xbp1s* was not rhythmic in *Bmal1-luc* NIH3T3 cells (Fig. S6C). These data suggest that 12hr RNA rhythms in NIH3T3 cells are driven by a different mechanism and primarily regulated by *Bmal1* itself or the circadian core clock machinery.

We next focused on the 161 12hr rhythmic RNAs and assessed which RNA process contributes to drive 12 hr RNA rhythms. By using adjusted B.H.q value, we found that only 12% (or 19/161 RNAs) of the 12 hr RNA rhythms had rhythmic RNA synthesis, whereas 67% (or 108/161 RNA) had rhythmic RNA degradation (Fig. 7A-B, Datafile 7). 12 hr RNA synthesis rhythms were less robust than 24 hr synthesis rhythms (Fig. 7C), while the relative amplitude of RNA processing and degradation rhythms were comparable between 24 hr and 12 hr (Fig. 7D). In addition, 12 hr RNA synthesis rhythms were less robust than 12 hr degradation rhythms (mean B.H.q-value: 0.804 (synthesis) vs 0.239 (degradation), p < 2.20 x 10^-16^, Mann-Whitney U-test) (Fig. 7C) and the relative amplitude of 12 hr degradation rhythms were higher than that of synthesis rhythm (mean relative amplitude: 0.045 (synthesis) vs 0.061 (degradation), p = 2.26 x 10^-7^, two-tailed Student’s t-test) (Fig. 7D). Unlike 24 hr RNA rhythms, there was no discernable phase relationship between synthesis, mature RNA level and degradation for 12 hr rhythms (Fig. 7E-F). Interestingly, the relative amplitude of 12hr RNA rhythms correlated with that of 12hr RNA degradation (Fig. 7H), but not with that of 12 hr RNA synthesis (Fig. 7G). These data collectively suggest that in NIH3T3 cells, 12 hr ultradian RNA rhythms are primarily driven by rhythmic degradation and regulated by *Bmal1* or the circadian core clock machinery.

**Figure 7.**
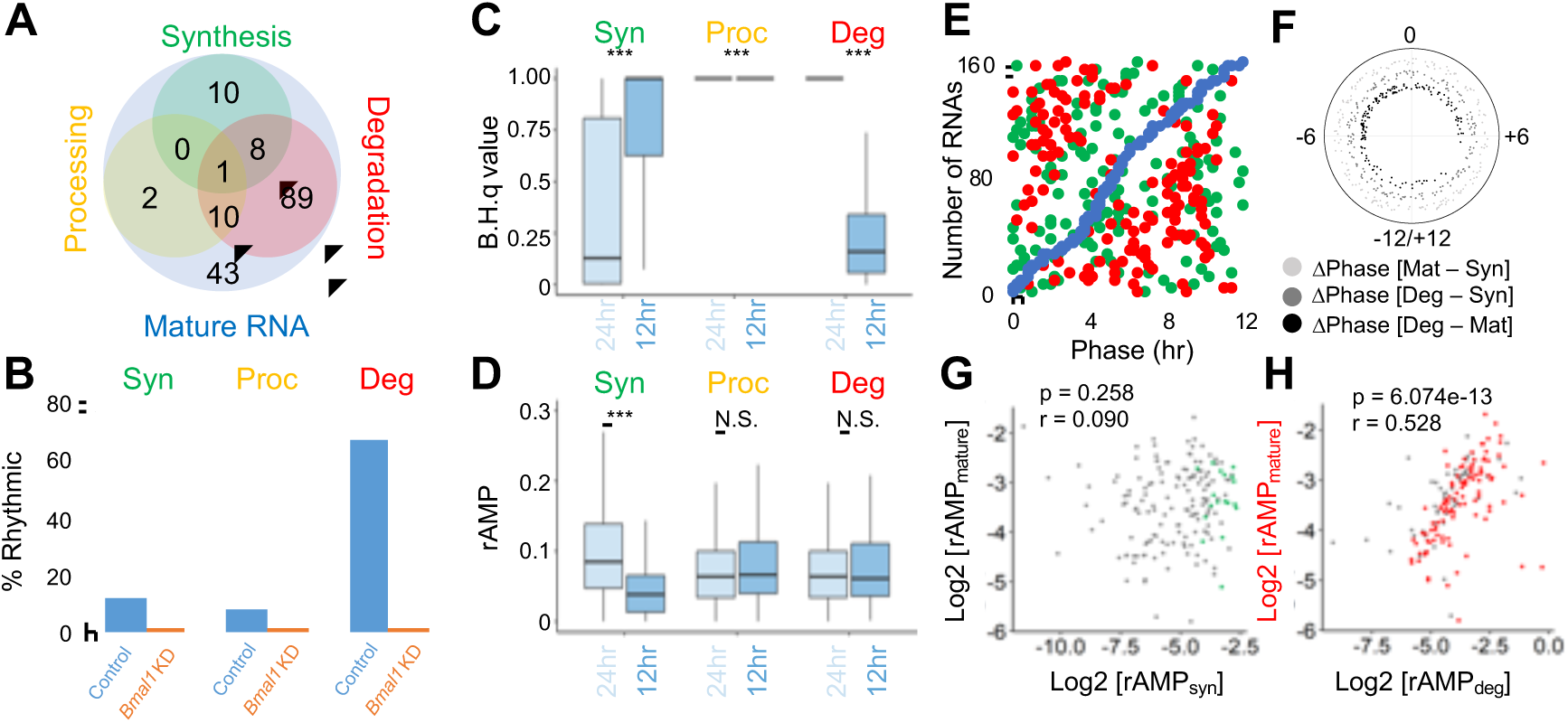
12hr RNA rhythms are primarily driven by rhythmic degradation. 161 rhythmic mature RNAs with a period of 12 hrs in control cells were analyzed. (A) Venn diagram showing the number of RNAs whose RNA synthesis (green), processing (yellow), and degradation (red) was rhythmic (metaCycle_BH.q < 0.25). (B) The percentage of RNAs with statistically significant synthesis, processing, or degradation rhythms (Meta2d BH.q < 0.25) among the 161 rhythmic mature RNAs with a period of 12 hrs in control (blue) and *Bmal1* (orange) KD cells. (C) Distribution of adjusted B.H.q values for the 685 24hr (left) or 161 12hr rhythmic RNAs in control cells. (D) Distribution of relative amplitude (meta2d_rAMP) for the 685 24hr (left) or 161 12hr rhythmic RNAs in control cells. Box plots represents two quartiles ± 1.5 interquartile range from median (midline). ***; p-value < 0.001, N.S.; p > 0.05 (C: two-tailed Mann-Whitney U-test, D: two-tailed Student’s t-test). (E) Peak phases of RNA synthesis rate (green), mature RNA levels (blue) and degradation rate (red). (F) Phase difference (hr) between mature RNA levels and RNA synthesis (light grey), mature RNA and RNA degradation (dark grey), and RNA synthesis and degradation (black). (G, H) Correlation between the relative amplitude of mature RNA and RNA synthesis rate (G) or degradation rate (H). All values are log2 transformed. r: Pearson correlation coefficient. Colored dots represent those that are statistically rhythmic (meta2d_B.H. q-value <0.25) hwhereas grey dots are statistically not rhythmic.

## Discussion

In this study, we used a metabolic labeling approach and monitored RNA dynamics in NIH3T3 cells around the circadian cycle. We found that a) rhythmic RNA synthesis is the major contributor in driving ∼24 hr rhythmic RNA expression (Fig. 2, 3), whereas rhythmic RNA degradation is important for 12hr RNA rhythms (Fig. 7), b) these rhythms were largely abolished in *Bmal1* KD cells (Fig. 2, 3, S6), and c) rhythmic expression of core clock genes are higher in amplitude and regulated by multiple rhythmic processes (Fig. 4K-L, 5). One of the limitations of our study is that we only had one biological replica (each taken from a single distinct cell culture dish) and samples were taken over one circadian cycle, reducing the statistical power of our analysis to some extent. We also used *in vitro* cell culture system, as metabolic labeling methodology is not currently applicable *in vivo*, and detected lower number of rhythmic RNAs compared to mouse tissues^9, 41^. Nevertheless, our data from NIH3T3 cells closely align with other circadian transcriptome and 4sU-seq studies^38, 47^ and provide critical RNA dynamics information to fully understand how the core circadian clock machinery shapes RNA rhythms.

Previous studies have assessed the effect of RNA synthesis (i.e., transcription) in regulating rhythmic RNA expression. In some studies, the level of pre-RNA (i.e., intronic levels in RNA-seq) was used as a proxy for RNA synthesis ^5, 13–15, 53^, while newer studies inferred RNA synthesis from directly measuring the level of nascent or actively transcribed RNAs ^13, 54, 55^. We therefore compared the RNA synthesis rate with the levels of newly synthesized RNAs in our dataset. As INSPEcT calculates RNA synthesis rate from the exonic levels in 4sU-seq ^37^, these two variables were linearly correlated (r = 1) (Fig. S7A). There was also a very strong correlation between RNA synthesis rate and the levels of newly synthesized RNAs (r = 0.964 with 4sU-seq [RPKM_gene_] and r = 0.837 with 4sU-seq [TPM_gene_]) (Fig. S7B-C). In support of this, the phase of RNA synthesis and that of the newly synthesized RNA levels aligned well with each other (fig. S7G) and the variation of their phase difference was small (²Phase_(Syn-4sUseq[RPKMgene]_ = 0.38 ± 1.54 hr , ²Phase_(Syn-4sUseq[TPMgene])_ = 0.03 ± 3.07 hr) (Fig. S7H). The correlation was slightly weaker between RNA synthesis rate and the pre-RNA level (i.e., intronic levels from RNA-seq) (r = 0.717) (Fig. S7D), and the phase difference was more variable (τιPhase_(Syn-RNAseq[RPKMintron])_: = -0.17 ± 4.57 hr) (Fig. S7G-H). The pre-RNA level also correlated strongly to that of newly synthesized RNAs (r = 0.765 with 4sU-seq [RPKM_gene_] and r = 0.810 with 4sU-seq [TPM_gene_]), but at a lesser degree compared to RNA synthesis rate (Fig. S7E-F). The difference in peak phase was also more variable between the levels of pre-RNA and newly synthesized RNAs (τιPhase_(PreRNA-4sUseq[RPKMgene]_ = 0.17 ± 4.41 hr and τιPhase_(PreRNA-4sUseq[TPMgene])_ = 0.12 ± 4.27 hr) (Fig. S7G-H). In fact, the correlation between pre-RNA and mature RNA was stronger (r > 0.96) and their phase difference was less than that between pre-RNA and RNA synthesis (τιPhase_(PreRNA-MatureRNA)_ = -0.03 ± 4.01 hr) (Fig. S2, Fig. S7G-H). These data collectively demonstrate that the RNA synthesis rate correlates stronger to the level of newly synthesized RNAs (i.e., 4sU-seq [RPKM_gene_] and 4sU-seq [TPM_gene_]) compared to the pre-RNA levels.

One of the surprising results was that rhythmic RNA degradation only had a modest contribution to 24 hr RNA rhythms at least in NIH3T3 cells (Fig. 2, 3). Even though several mechanisms, such as miRNA, RNA-binding proteins, and poly(A) tail length, have been implicated in regulating rhythmic RNA degradation ^3^, our data indicate that these mechanisms don’t have a strong impact, or are not rhythmically regulated at least in NIH3T3 cells. Interestingly, however, RNA degradation appeared to be weakly rhythmic (Fig. 3: top right), and this rhythmicity was abolished in *Bmal1* KD cells (Fig. 3: bottom right). How can rhythmic degradation, albeit weak, be controlled by *Bmal1* or the core clock machinery? Because the amplitude of RNA degradation rhythms is generally much lower than those of RNA synthesis (Fig. 3D) and we don’t see the global effect of rhythmic degradation (Fig. 2), we propose that the weak RNA degradation rhythm is regulated by a transcript-specific manner, and potentially coupled with rhythmic RNA synthesis. One possible mechanism is transcript buffering, a cellular mechanism to maintain the homeostasis of transcript level. The rates of RNA synthesis and degradation can be kinetically coupled by establishing a feedback loop between RNA synthesis in the nucleus to degradation of their corresponding RNAs in the cytoplasm^56–63^. This mechanism has been well established in yeast and recently demonstrated in human cells ^64, 65^ however, all the studies were done in a steady-state condition and its regulatory mechanisms, particularly under a dynamic condition, remain elusive (reviewed in ^57^). Another possibility is the rhythms in poly(A) tail length, as has also been suggested to be coupled with rhythmic RNA synthesis ^19^. Indeed, several rhythmic mature RNAs detected in our dataset exhibited rhythmic poly(A) tail length in mouse liver ^19^. It will be of future interest to uncouple RNA synthesis from degradation, and test whether degradation is still rhythmic even in the absence of rhythmic RNA synthesis.

Our study also highlighted the robust control of rhythmic core clock RNA expression (Fig. 4, 5). Given that core clock genes are rhythmic in many more tissues compared to clock-controlled genes ^9, 10^, it is reasonable that their RNA rhythmicity is regulated by multiple steps to ensure that cell-autonomous rhythmicity with high amplitude can be sustained even in the presence of environmental noise. What makes their amplitude high? Because circadian cistrome datasets for various core clock proteins as well as different forms of RNA polymerase II are already available in mouse liver ^14, 53, 66^, analyses of their dynamics will likely provide an important clue as to what drives the high amplitude of core clock gene expression.

Overall, our study further elucidates the extent by which transcriptional and post-transcriptional regulatory steps drive rhythmic RNA expression. It would be of great interest to expand our study and explore the dynamics of rhythmic protein expression in the future to fully understand how the core circadian clock machinery regulates gene expression rhythms and ultimately rhythmic downstream processes.

## Materials and Methods

### Cell culture and lentiviral transduction

NIH3T3 derived *Bmal1-luc*^31^ and HEK293/T17 cells were cultured with Dulbecco’s Modified Eagle Medium (DMEM) (Gibco: Cat #11965118) supplemented with 10 % fetal bovine serum (Corning, Cat#MT35010CV) at 37°C with 5.0 % CO2. We followed a standard procedure to prepare lentiviruses containing short-hairpin (sh) RNAs ^67^. Briefly, viral particles were prepared in HEK293/T17 cells by transfecting pLP-1, pLP-2, and pLP-VSVG plasmids (ThermoFisher) along with pLL3.7GW vector containing shRNAs for control (target sequence: CAACAAGATGAAGAAGAG CACCA) or *Bmal1* (target sequence: GGAAGGATCAAGAATGCAA) using polyethylenimine (Polysciences Inc., #23966) ^68^. 48 hrs after the DNA transfection, culture medium containing viral particles were collected and filtered by 0.22 µm PDVF membrane (Millipore, Cat#SLGV013SL). The media was further ultracentrifuged at 75,000g for 2 hrs at 10°C before being reconstituted to the same volume of fresh cell culture media. *Bmal1-luc* NIH3T3 cells were then transduced with polybrene (Milipore, Cat#TR-1003-G), and media containing viruses were replaced with fresh media 24 hrs after transduction. Transfection and transduction efficiency was visually inspected by EGFP signals that is encoded in the pLL3.7GW vector.

### 4sU labeling and RNA pull-down

72 hrs after the viral transduction, *Bmal1-luc* NIH3T3 cells were labeled with 400μM of 4-thiouridine (4sU) (Cayman Chemical, Cat#16373) for 2 hrs. Subsequently, RNAs were extracted with TRIzol (Invitrogen, Cat# 15596018) according to manufacturer’s instructions, and then treated with DNase I (ThermoFisher, AM2239) at 37°C for 1 hr. After re-purifying RNAs using TRIzol, 1 µg of yeast spike-in RNA (gift from Dr. Silke Hauf: Virginia Tech) labeled with 4-thiouracil (Neta Scientific, CMX-21484-1G) for 1 hr, and 300 fmol 4sU-labeled DNA/RNA hybrid spike-in control (ATTTAGGTGACACTATAGGATCCTCTAGAGTCGACCTTCTCCCTATAGTGAGTC GTATTAGCA[4sU]CAG) were added to each RNA sample to normalize the RNA pull-down efficiency (see below). 4sU-labeled RNA was then biotinylated with 1mg/ml EZ-link HPDP-Biotin (ThermoFisher, cat: 21341) for 90 min at room temperature as previously described^34, 69, 70^. 30ug RNA (approximately 50% for sh*Ctrl* and 60-70% for sh*Bmal1* samples) were subsequently subjected to streptavidin pull-down, while the remainder was set aside as ‘total RNA’. Streptavidin magnetic beads (ThermoFisher cat: 65001) were pre-washed with 0.1M NaOH, 0.1M NaCl, then washing buffer (10mM Tris-HCl pH 7.4, 10mM EDTA, 1M NaCl) to remove RNase. Beads were then incubated with RNA samples in Washing Buffer (10 mM Tris-HCl pH7.4, 1M NaCl, 10 mM EDTA) for 45 min at room temperature with rotation. Beads were washed twice with washing buffer for 5 min and all the flow through (‘pre-existing’) fractions from each wash were separated from the bead bound fraction using a DynaMag-2 magnetic rack (Life Technologies, Cat#12321D) and pooled together. After the second wash, RNAs bound to the beads were recovered first by incubating the beads with 0.1 M DTT for 10 minutes at room temperature. The remaining beads were further incubated with pre-warmed 0.1 M DTT at 65 °C for 10 min and all the RNAs were collected in a single tube (‘newly synthesized’). RNAs from both pre-existing and newly synthesized fractions were precipitated with ethanol with 1M NaCl and 20 µg glycogen (Ambion; Cat#9510) and reconstituted in Diethyl pyrocabonate (DEPC) treated deionized and distilled water (DDW).

### RNA pull-down efficiency calculation

We calculated the pull-down efficiency using 4sU-labeled DNA/RNA hybrid spike-in control, as was previously described^34^. Briefly, RNAs from both pull-down and flow-through fractions were purified by TRIzol and responded in the same volume of DEPC treated DDW. An equal volume of RNA solution from both fractions was used to generate cDNA separately using High-Capacity cDNA Reverse Transcriptase Kit (Applied Biosystems, REF4368813) according to the manufacturers’ instruction. We quantified the relative amount of cDNA in each fraction using QuantStudio6 (Life Tech) with PowerSYBR Green PCR Master Mix (Applied Biosystems, REF4367659). We determined the pull-down efficiency by calculating the ratio of the level of spike-in control between the pull-down fraction and the combination of pull-down and flow through fractions).

### Transcriptomic analyses

Approximately 100-200ng of total and newly synthesized RNAs from each time point were subjected to transcriptomic analysis (Azenta Biosciences). RNA quality of each sample was verified by the RNA integrity number (RIN) > 9.5 measured by Tapestation 4200 (Agilent). Additional DNase I treatment was performed before proceeding to the library preparation per recommendation from Azenta. Paired-end, strand-specific, and rRNA depleted sequencing library was then prepared using QIAGEN FastSelect rRNA HMR Kit (Qiagen, cat#334378) and NEB Directional Ultra II RNA library prep kit (NEB, E7760) by following the manufacturer’s recommendations (NEB, cat# E7765). Qubit 2.0 Fluorometer (ThermoFisher Scientific) and quantitative PCR (KAPA Biosystems) method were used to quantify each library before multiplexing and the distribution of reads lengths were measured by the Agilent Tapestation 4200 (Agilent Technologies). Libraries were then sequenced by Illumina Hi-seq.

### Bioinformatical analyses

Azenta used their in-house quality control bioinformatical pipeline to return FASTQ files with a quality score of >Q30 on average. We further used TrimGalore (Martin et al. 2010) to remove adapter sequences and remove reads of low quality (<Q20), which we found to be about 0.2% per file. Reads were mapped with STARv2.7.7a (Dobin et al. 2013) using options -- outFilterScoreMinOverLread 0.3 and --sjdbOverhang 100. Reads mapped to the NR003278.3 (*Mus musculus* 18S rRNA) and NR003279.1 (*Mus musculus* 28S rRNA) were eliminated first. Remaining reads were then mapped to the mouse mm10 genome (GENECODE: GRCm38.p6.genome.fa) and read counts against gene (transcript per million or TPM), exon, and intron were independently quantified with HOMER (V4.11.1) (Heinz et al. 2010) using gencode.vM25.annotation.gtf. using option condenseGenes. Transcripts with TPM < 0.25 (as an average of all time-points and conditions) were removed for all downstream analysis. Reads per kilo base per million mapped (RPKM) was also calculated using HOMER (v4.11) to quantify the reads mapped to exon or intron separately with the –exon or –intron option for both RNA-seq and 4sU-seq. By using R (V4.1.0), we supplied these four input data files to INSPEcT (V1.22.0) ^37^ using the ratesFirstGuess options: 2 hr labeling time, deg=FALSE to obtain a total of five output files. INSPEcT eliminates single-exon genes as it requires both exonic and intronic levels to return these output files.

### Other computational analyses

#### Read count distribution

Read counts for the intron and exon were from HOMER (v4.11) after mapping raw reads to the mouse transcripts (GRCm38.p6, gencode.vM25.annotation.gtf) using STAR (v2.7.7a). Unmapped reads were then mapped to the mouse genome (GRCm38.p6, gencode.vM25.annotation.gtf) using STAR (v2.7.7a) and the reads uniquely mapping to the genome but not the transcript sequences were determined to be intergenic. The average of all time points within each dataset were then calculated.

#### Rhythmicity analysis

We used MetaCycle (v1.2.0, CRAN R package) ^40^ and used meta2d B.H. q-values to define rhythmicity. To detect rhythmicity with different periods, we set the period range to 20-28, 10-14, and 6.7-9.3, for 24hr, 12hr, and 8 hr periods, respectively. These ranges reflect a 16.6% variation of the period.

#### Sequence data visualization (GBiB)

The bam files generated by STAR (see above) were sorted and indexed by Samtools (version 1.11) ^71^, and converted into bigWig files by DeepTools (version 3.5.0) ^72^ with the option -filterRNAstrand in order to visualize strand specifically. The files were uploaded to UCSC Genome Browser using the Genome Browser in a Box (GBiB)^73^ to visualize mapped reads.

#### Phase-sorted heatmaps

Mature RNA expression data were first log_2_ transformed and Z-scores for each RNA profile were manually calculated. The data were then sorted by meta2d_phase and heatmaps were generated by gplots heatmap.2 function ^74^ using Rstudio (v4.1.0) with no additional scaling options.

#### PCA analysis

Principle component analysis was performed using tidyverse prcomp function in R (1.3.2) ^75^ with scaling = TRUE option.

### Statistical analyses

The statistical tests were performed using Rstudio (v4.1.0). The default R “stats” package v4.1.0 was used for Pearson correlation test, Student’s t.test and Mann-Whitney U-test, while the “car” package (v3.1) was used for the Levene test. The “circular” package was used for the Watson-Wheeler test (v0.4-95) with Rstudio(v.4.2.1).

## Acknowledgments

Bmal1-luc NIH3T3 cells^31^ is a gift from Dr. Ueli Schibler (University of Geneva). Both control and *Bmal1* knock-down plasmids ^68^ were a gift from Dr. Andrew Liu (University of Florida). Yeast spike-in was a gift from Dr. Silke Hauf (Virginia Tech). Dr. Mattia Furlan (Istituto Italiano di Technologia) provided technical assistance in running the INSPEcT algorithm ^37^ and Evan S. Littleton, and Lin Miao also provided other technical assistance. This work was supported by National Institute of Health F31 AG071393 (to B.A.U) and R01 GM126223 (to S.K.).

## Data availability

Both 4sU-seq and RNA-seq data will be made available upon publication. They will be deposited to NIH SRA (Submission ID: 13164610).

## Author contributions

B.A.U. and D.E.W acquired and analyzed the data. S.K. conceived and designed the project, acquired funds, and wrote the manuscript.

