## Supplemental Information for "Coordination of rhythmic RNA synthesis and degradation orchestrates 24-hour and 12-hour RNA expression patterns in mouse fibroblasts"

**Supplementary methods:**

*Real-time bioluminescence recordings*

After cells were split into 35-mm dishes became confluent, the medium was replaced with DMEM supplemented with 50% horse serum for 2 hr. After synchronizing cells, medium was then changed to phenol red-free DMEM (Cellgro, 90-013-PB) supplemented with 100 µM luciferin, 10 mM HEPES (pH 7.2), 1 mM sodium pyruvate, 0.035% sodium bicarbonate, 2% FBS, 1x penicillin/streptomycin, and 2 mM L-glutamine. Real-time bioluminescence recordings were performed using a LumiCycle (Actimetrics, Inc.). 24 hr after serum shock, a dish was removed from the LumiCycle and 400µM of 4-thiouridine (4sU) was added (Cayman Chemical, Cat#16373). The 35mm dish was then immediately placed back into the LumiCycle to continue recording. This process was repeated every 4 hrs for 24 hrs, each plate only being labeled once (n=2).

*Comparison between datasets*

To analyze the microarray data, we used GCRMA (version 2.66.0) in R, with the Mouse Genome 430 2.0 or U74A (downloaded from Bioconductor mouse4302.db or pd.mg.u74av2) annotations with GCRMA quantile normalization and background normalization. RNA-seq data [77] were mapped to the mouse mm10 genome (UCSC: mm10.fa, mm10.ncbiRefSeq.gtf) with STAR (v2.7.7a) and quantified with HOMER (V4.11.1) to calculate TPM with the option condenseGenes. Data points between 0-20[77], 0-48[76]. or 24- 47(1 h) or 24-46(2 h)[41] were used. We selected the 11313 genes used in our dataset from each dataset to make them comparable.

Supplemental Figure 1: Unruh et al.,

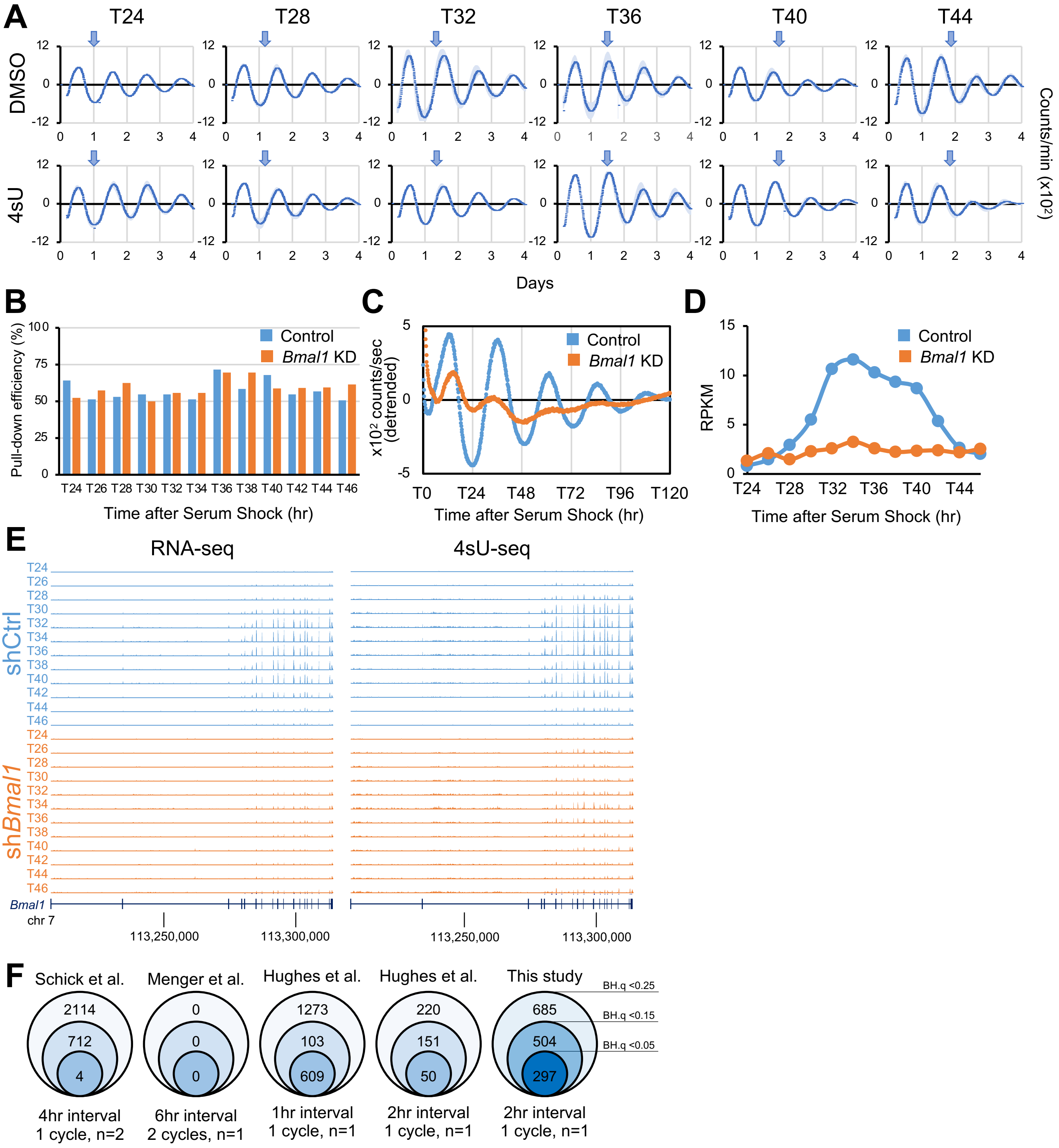


**Figure S1. Validation of experimental conditions for rhythmic 4sU-seq and RNA-seq analysis.** (A) *Bmal1-luc* bioluminescent output with a mock "pulse-in" procedure. DMSO (top) or 400 μM 4sU (bottom) was applied at each time point indicated by blue arrows. The data represent mean ± SEM (n=2). (B) Pull-down efficiency of 4sU-lableled DNA/RNA hybrid spike-in control (n=1). (C-D) Verification of the *Bmal1* knock-down by monitoring bioluminescence output (n=1) (C) or *Bmal1* mature RNA levels from the RNA-seq analysis (n=1) (D). (E) Genome browser view of mouse chromosome 7 at *Bmal1* locus for total (left) or newly synthesized (right) RNA fractions from both control (blue) and *Bmal1* KD (orange) cells. (F) Number of rhythmic genes detected in NIH3T3 transcriptome datasets with various experimental conditions. Only the 11313 genes included in our dataset were considered in each dataset. Note that Schick dataset is from RNA-seq while Menger and Hughes datasets are from microarray. Schick datasets were collected T0-T24, while other datasets were collected T24-T48. We performed in-house MetaCycle analysis for all the data.

Supplemental Figure 2: Unruh et al.,

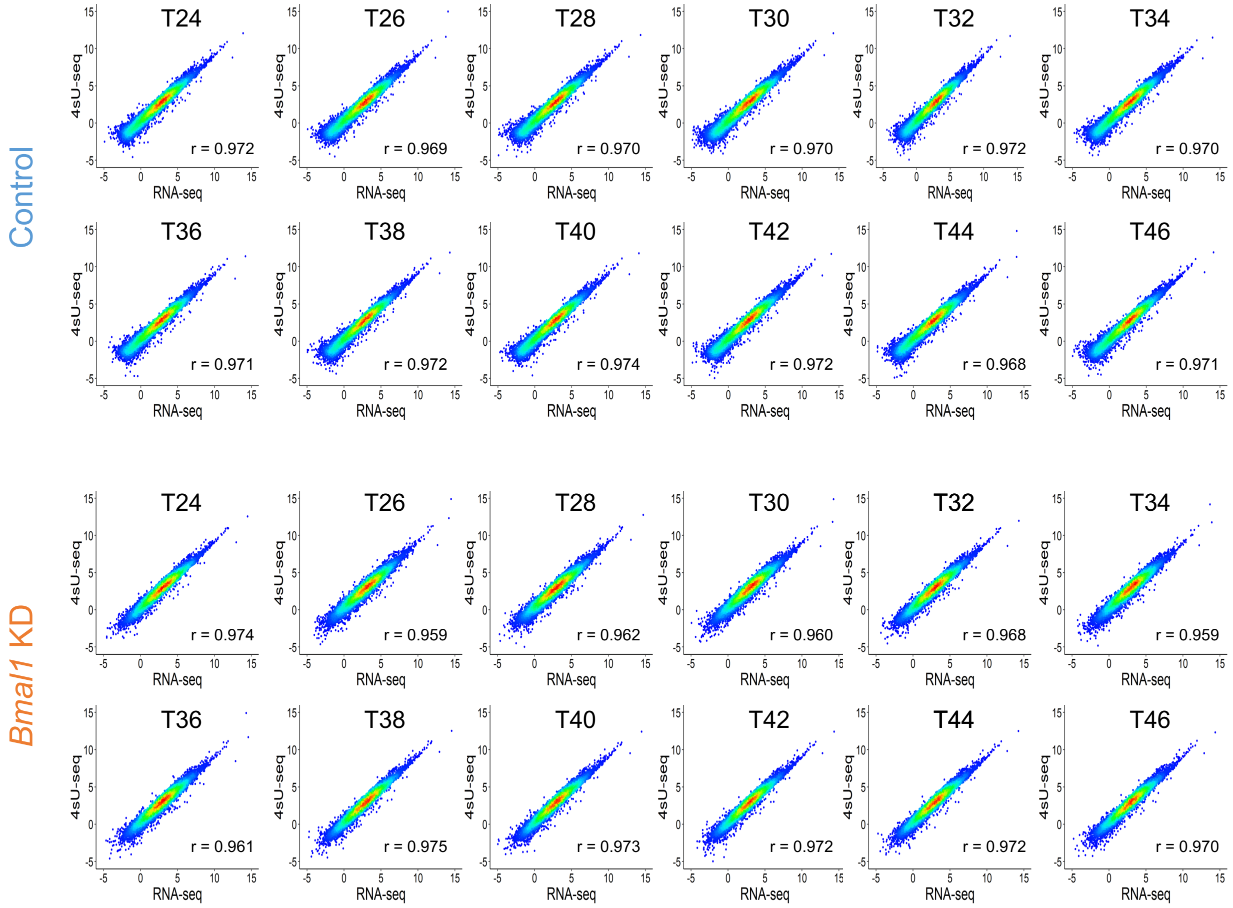


**Figure S2: Correlation of RPKM exon reads between RNA-seq and 4sU-seq at each time point.** X axis represents log2(RPKM-exon) from RNA-seq while y axis represents log2(RPKM-exon) from 4sU-seq. r: Pearson correlation coefficient.

Supplemental Figure 3: Unruh et al.,

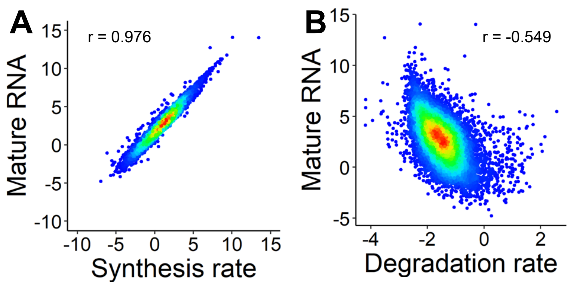


**Figure S3: Correlation between the level of mature RNAs and log2 transformed rates of synthesis or degradation.** Average of all time points was used. r: Pearson correlation coefficient.

Supplemental Figure 4: Unruh et al.,

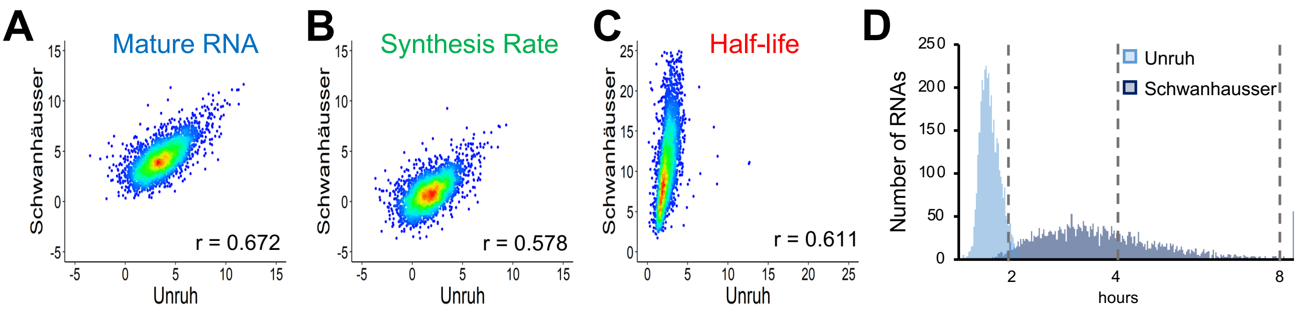


**Figure S4: Comparison between this study (Unruh) and a previous study including both RNA-seq and 4sU-seq at a steady-state (Schwanhäusser et al., 2011).** (A) Mature RNA levels (log2(RPKM-Total$_{exon}$), (B) Log2 transformed synthesis rate, (C) RNA half-life (hr). Only genes included in both studies were compared (n= 3534). r: Pearson correlation coefficient. (D) Distribution of RNA half-life between this study (median: 2.27 hr) and Schwanhausser (median: 10.04 hr).

Supplemental Figure 5: Unruh et al.,

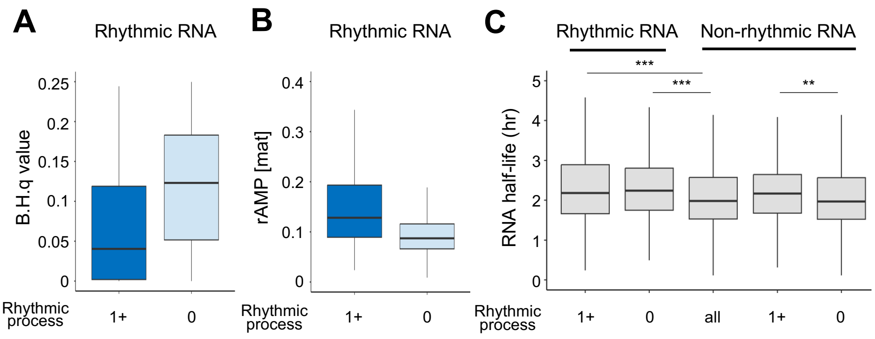


**Figure S5. Characteristics of rhythmic and non-rhythmic RNAs.** (A) Distribution of meta2d_B.H.q values for the 685 rhythmic RNAs . $p < 2.2 \times 10^{-16}$ (two tailed Mann-Whiney U test) between those that have one or more rhythmic process vs without any rhythmic process. (B) Distribution of relative amplitude (rAMP) for the 685 rhythmic RNAs. $p < 2.2 \times 10^{-16}$ (two-tailed Student's t-test) between those that have one or more rhythmic process vs without any rhythmic process. (C) RNA half-lives of rhythmic and non-rhythmic RNAs, depending on whether one or more process is also rhythmic. ***; $p < 0.001$, **; $p < 0.01$, (two-tailed Student's t-test). Box plots represents two quartiles ± 1.5 interquartile range from median (midline).

Supplemental Figure 6: Unruh et al.,

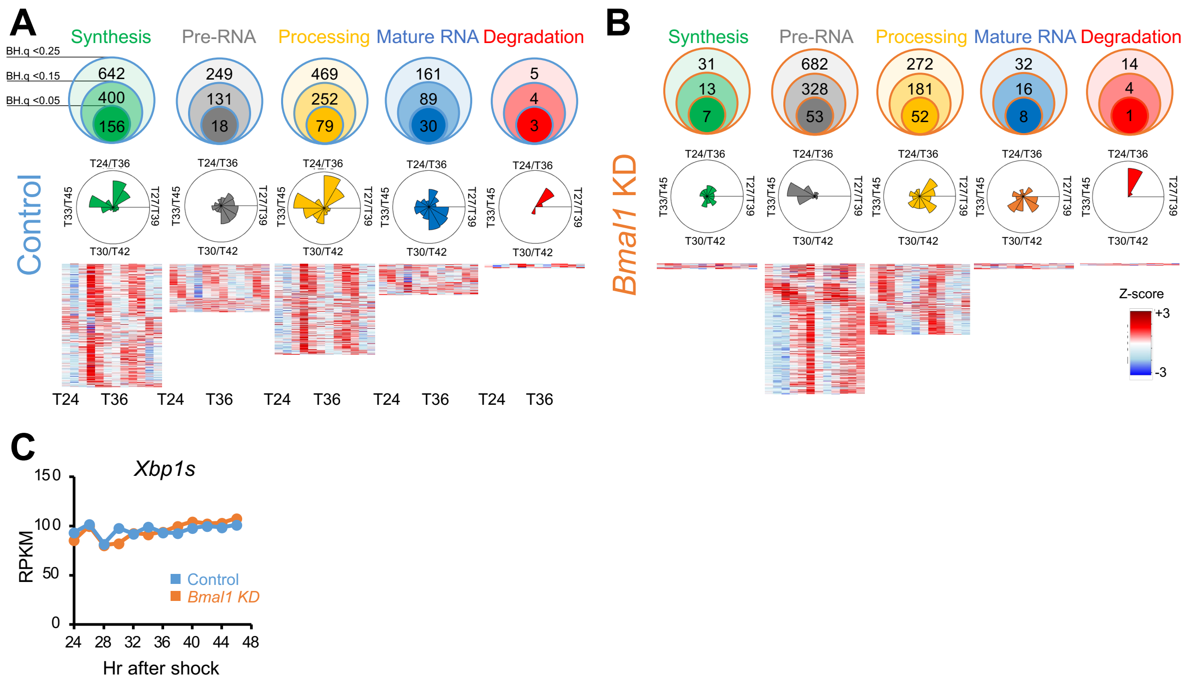


**Figure S6. 12 hr rhythms are also regulated by *Bmal1* or the core clock machinery.** A total of 11313 genes were analyzed in all datasets. (A, B: top) Number of RNAs whose RNA synthesis (green), processing (yellow), or degradation (red) as well as unprocessed (grey) or mature RNA (blue) levels were ~12 hr rhythmic with various statistical thresholds (meta2d_B.H.q < 0.25, 0.15, or 0.05) in control (A) or *Bmal1* KD (B) cells. (A, B: middle) Circular histograms representing the peak phase distribution of ~12 hr rhythmic RNA synthesis (green), processing (yellow), or degradation as well as pre- (grey) or mature RNA (blue) levels in control (A) or *Bmal1* KD (B) cells. Each bin represents 2 hrs. Radius line represents number of genes at each tick mark, and the most outer line is 200 (synthesis), 70 (pre-RNA), 100 (processing), 35 (mature RNA), and 5 (degradation) RNAs in control and 10 (synthesis), 300 (pre-RNA), 35 (processing), 10 (mature RNA), and 15 (degradation) RNAs in *Bmal1* KD cells. (A, B: bottom) Phase-sorted heatmap of ~12 hr rhythmic RNA synthesis (green), pre-RNA levels (grey), processing (yellow), mature RNAs (blue), and degradation (red) in control (A) or *Bmal1* KD (B) cells. Color represents Z-scores, in which red is high while blue is low. Each line represents one RNA. (C) RNA expression level of *Xbp1s* (RPKM) in control (blue) and *Bmal1* KD (orange) cells.

Supplemental Figure 7: Unruh et al.,

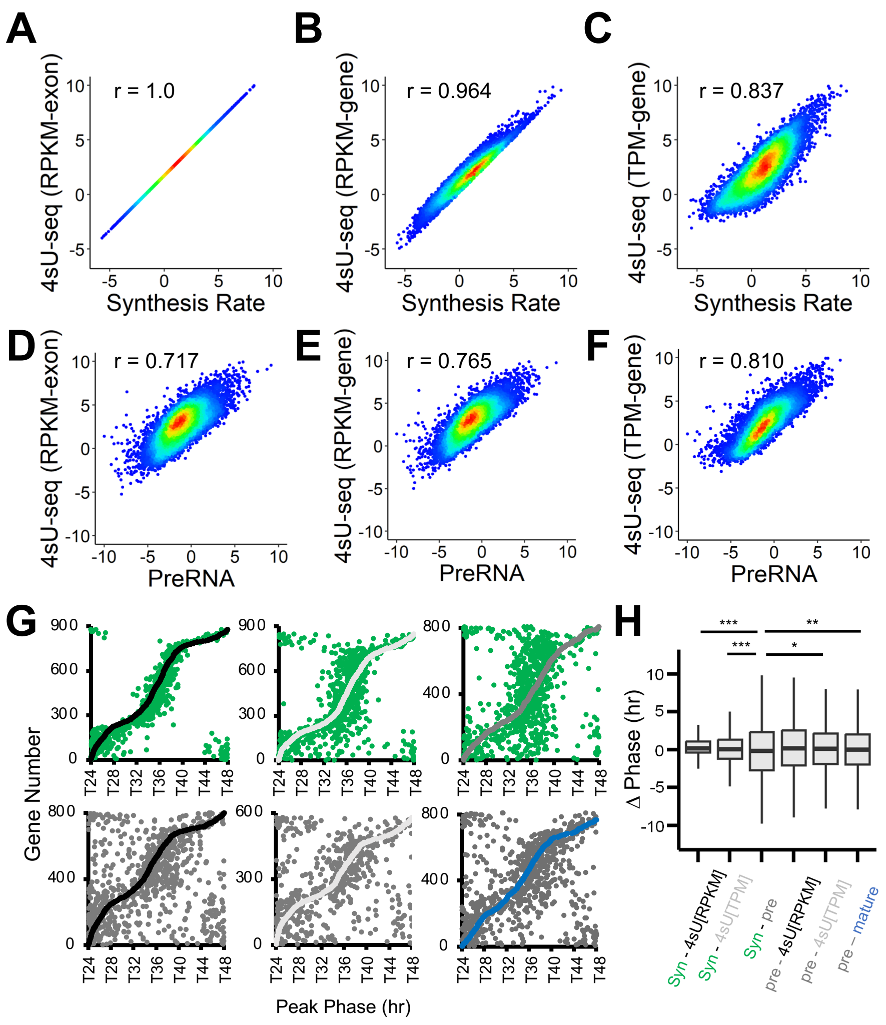


**Figure S7: Comparison between RNA synthesis rate or pre-RNA levels and various quantification methods from RNA-seq and 4sU-seq.** (A) RNA synthesis rate and 4sU-seq [$RPKM_{exon}$], (B) RNA synthesis rate and 4sU-seq [$RPKM_{gene}$], (C) RNA synthesis rate and 4sU-seq [$TPM_{gene}$], (D) pre-RNA (or RNA-seq [$RPKM_{intron}$]) and 4sU-seq [$RPKM_{exon}$], (E) pre-RNA and 4sU-seq [$RPKM_{gene}$], and (F) pre-RNA and 4sU-seq [$TPM_{gene}$]. The diagonal line in (B) represents genes in which the $RPKM_{exon}$ = $RPKM_{intron}$. Note that INSPEcT eliminates all genes when the RPKM of exons are lower than that of introns and the synthesis rate of these genes cannot be calculated. All data are log2 transformed and represent average of all time points data in control cells for all 11313 genes. r: Pearson correlation coefficient. (G) Peak phase relationship between RNA synthesis rate (green) and 4sU-seq [$RPKM_{gene}$] (black) (top left), RNA synthesis rate (green) and 4sU-seq [$TPM_{gene}$] (light grey) (top middle), RNA synthesis rate (green) and pre-RNA (or 4sU-seq [$RPKM_{exon}$]) (grey) (top right), pre-RNA (grey) and 4sU-seq [$RPKM_{gene}$] (black) (bottom left), pre-RNA (grey) and 4sU-seq [$TPM_{gene}$] (light grey) (bottom middle), pre-RNA (grey) and mature RNA (blue) (bottom right). The peak phases were calculated as meta2d_phase and depicted are those with meta2d B.H.q < 0.25 in either measurements. (H) Variation of the phase difference shown in D. Boxplots represent two quartiles ± 1.5 interquartile range from median (midline) (n = 848: Syn-4sU[RPKM], 879: Syn-4sU[TPM], 807: Syn-pre, 798: pre-4sU[RPKM], 579: pre-4sU[TPM], and 809: pre-mature]). ***; p < 0.001, **; p < 0.01 *; p < 0.05 (Levene's test).
